# Potential short-term negative versus positive effects of olive mill-derived biochar on nutrient availability in a calcareous loamy sandy soil

**DOI:** 10.1101/2020.04.23.057695

**Authors:** Azzaz Alazzaz, Adel R. A. Usman, Munir Ahmad, Jamal Elfaki, Abdelazeem S. Sallam, Hesham Ibrahim, Mutair A. Akanji, Mohammad I. Al-Wabel

## Abstract

In this study, a greenhouse pot experiment with maize (*Zea mays L*.) was conducted using treatments consisting of a control (CK), inorganic fertilizer of NPK (INF), and 1% and 3% (wt/wt) of olive mill solid waste (OMSW)-derived biochar (BC) at various pyrolytic temperatures (300–700 °C). The goal was to investigate potential negative versus positive effects of BC on pH, electrical conductivity (EC), and nutrient (P, K, Na, Ca, Mg, Fe, Mn, Zn, and Cu) availability in a calcareous loamy sandy soil. The results showed that application of OMSW-derived BC, especially with increasing pyrolysis temperature and/or application rate, significantly increased soil pH, EC, NH_4_OAc-extractable K, Na, Ca, and Mg, and ammonium bicarbonate-diethylenetriaminepentaacetic acid (AB-DTPA)-extractable Fe and Zn, while AB-DTPA-extractable Mn decreased. The application of 1% and 3% BC, respectively, increased the NH4OAc-extractable K by 2.5 and 5.2-fold for BC300, by 3.2 and 8.0-fold for BC500, and by 3.3 and 8.9-fold for BC700 compared with that of untreated soil. The results also showed significant increases in shoot content of K, Na, and Zn, while there were significant decreases in shoot content of P, Ca, Mg, and Mn. Furthermore, no significant effects were observed for growth of maize plants as a result of biochar BC addition. In conclusion, OMSW-derived BC can potentially have positive effects on the enhancement of soil K availability and its plant content but it reduced shoot nutrients, specifically P, Ca, Mg, and Mn; therefore, application of OMSW-derived BC to calcareous soil might be restricted.

## Introduction

Rapid expansion in agriculture to feed the continuously growing world population has increased conventional intensification in farming systems, which consequently has resulted in soil nutrient depletion and various environmental concerns [1]. Intensive application of agro-chemicals has resulted in land degradation and declining soil health and quality [2, 3]. Hence, a global transition towards modern farming systems with sustainable soil health, safe ecosystems, food and nutrition security, and climate change mitigation is required. In this context, organic soil additives can potentially improve crop production, soil organic matter, rehabilitation of degraded land, and microbial activity with minimal environmental damage [4–7]. Among soil additives, biochar has been recently suggested as an emerging organic conditioner that can aid in overcoming soil problems and enhancing soil productivity. It can also be used as a tool for carbon sequestration to mitigate climate [8–10].

Biochar is produced from organic materials (e.g., biomass) through pyrolytic processes under limited oxygen and is generally characterized by its high content of fixed carbon [11]. Biochar materials are largely employed as additives to overcome soil problems and limitations by enhancing soil properties in relation to chemical, biological, and hydro-physical parameters, as well as nutrient content and efficiency [8, 10, 12–14]. These improvements in soil properties are mainly the result of positive characteristics of biochar, making it an ideal candidate to improve soils for sustainable agriculture. However, most investigations regarding biochar benefits have been conducted in acidic soils. Previous studies have demonstrated that biochar enhances the properties of acidic soils, mainly because of the liming effect induced by the alkaline nature of biochar [15, 16]. However, the effects of biochar application on the properties of alkaline soils with high pH and CaCO_3_ values are not well understood and have received much less attention. Previous studies have reported contrasting effects of biochar addition on soil and plants in alkaline soil [17–19].

Both the source of biomass and the pyrolysis conditions are considered the main factors determining the biochar properties, and thus, the effects of its application in the environment [20, 21]. On the other hand, selection of suitable and low-cost feedstock is of critical importance. Various types of feedstock, such as wood, grass, manure, sludge, paper, and industrial organic wastes, have been used for biochar production [11, 21, 22]. Among the various wastes generated in the environment, solid olive-derived waste might be applied as a potential low-cost feedstock that can be recycled into biochar.

The amount of olive mill solid waste (OMSW) produced globally accounted for 4 × 108 kg dry matter, comprising skin, pulp, and seeds, consisting of 38–50% (w/w) cellulose, 23–32% hemi-cellulose, and 15–25% lignin. The deposition of OMSW in the environment can have negative effects on water resources, soil microbes, and plants [23]. Physicochemical treatments can be used to recycle OMSW for the extraction of oil, fiber, polyphenols, pectins, or energy [24, 25] for its use as a soil amendment [26], compost [27], enzymes, fuel, or livestock feed [28]. Previous studies have demonstrated that the biochar produced from OMSW is a promising sorbent/immobilizing agent for removing heavy metals from aqueous solutions or to remediate heavy metal contaminated soils [26, 29]. Additionally, the enhancing effects of OMSW biochar on soil microbial biomass C and N, and alteration of the structure of the bacterial community has been reported by other authors [30, 31]. However, to date, there is no available information on the effects of OMSW-derived biochar on nutrient availability in alkaline sandy soils. Therefore, the aims of this study were to investigate (i) the effects of pyrolysis temperature (300–700 °C) on the properties of OMSW-derived biochar, (ii) potential negative versus positive effects of OMSW-derived biochar on the chemical properties and nutrients availability in calcareous loamy sandy soil and on maize (Zea mays L.) growth.

## Material and methods

### Production and characteristics of biochar

The solid waste from the olive mill was collected from the Al Jawf region, Saudi Arabia, and dried at 60 °C. The prepared OMSW feedstock was pyrolyzed in a closed container by furnaces under limited oxygen supply at temperatures 300, 400, 500, 600, and 700 °C for 4 h at 5 °C min^−1^. Biochar samples were collected, cooled in a desiccator, ground, sieved through a 2 mm sieve, labeled as BC300, BC400, BC500, BC600, or BC700, and stored for further analyses.

In OMSW-derived biochar (BC), the moisture content, and that of volatile and resident materials, and ash (proximate characteristics) were analyzed by the ASTM D1762-84 method [32]. The pH of OMSW-derived BCs was measured in a mixture (1:25, w/v) of BC and deionized water using digital pH meter. After measuring biochar pH, the mixture of BC and water was extracted to determine the electrical conductivity (EC) using a digital EC meter. Additionally, BC samples were analyzed with a scanning electron microscope (SEM; FEI, Inspect S50), X-ray diffraction (XRD; XRD-7000; Shimadzu Corp, Kyoto, Japan), surface area analyzer (ASAP 2020, Micromeritics, USA), and the Fourier transformation infrared method (Nicolet 6700 FTIR).

### Greenhouse pot experiment

The collected soil samples from an agricultural farm located in a dry land region (at Riyadh, Saudi Arabia) were air-dried, sieved, and analyzed for their physico-chemical properties. To identify particle size distribution according to Bouyoucos [33], the hydrometer method was applied. According to Sparks [34], and Nelson and Sommers [35], chemical soil properties, including pH, EC, CaCO3, and organic matter, were measured. The data for soil analyses showed that the soil samples, which were characterized as having a loamy sand texture (containing 80.89% sand, 12.07% silt, and 7.04% clay) and a low level of organic matter (0.12%), had an alkaline pH (measured in 1:1 suspension of soil to water) value of 7.8, EC (measured in 1:1 extracts of soil to water) value of 2.0 dS m-1, and high CaCO3 content (16.51%).

A greenhouse pot experiment with maize plants was conducted. Specifically, 1 kg of soil treated with 1% and 3% (wt/wt) of OMSW-derived BC at various pyrolytic temperatures (300 °C, BC300; 500 °C, BC500; and 700 °C, BC700). Additionally, treatments consisting of a control (CK) and inorganic fertilizer of NPK (INF) were applied in the study for comparison. Then, the treated and untreated soil samples were placed in pots, irrigated at the level of field capacity, and incubated for 21 d under laboratory conditions (at a temperature of 23 ± 2 °C). After the incubation period, three replications of the treated and untreated pots were transferred to the greenhouse and placed in a randomized complete block design. Ten maize seeds were sown in each pot. After seedlings emerged, the plants in each pot were thinned to four. The planting period lasted 4 weeks. During the growth period, the plants were irrigated and maintained at 70% of field capacity by weight loss. After 4 weeks of cultivation, the shoots of maize plants and soil were collected from the pots. The collected soil samples were air-dried and analyzed for pH, EC, AB-DTPA-extractable nutrients (P, Fe, Mn, Zn, and Cu), and ammonium acetate-(NH_4_OAc)-extractable K, Na, Ca, and Mg [34]. Additionally, the shoots of maize plants were collected, dried at 70 °C, and analyzed for P, Ca, Mg, K, Na, Fe, Mn, Zn, and Cu. The maize plant shoot samples were dried and digested using a dry ashing method [36]. In this study, inductively coupled plasma (ICP, Perkin Elmer Optima 4300 DV ICP-OES) was used to measure the concentrations of P, Ca, Mg, Fe, Mn, Zn, and Cu. The concentrations of K and Na were analyzed using a flame photometer.

### Statistics

The least significant difference (LSD) test (at the 0.05 level) using the Statistica software program was computed to compare treatment effects on the obtained parameters [37].

## Results and Discussions

### Characteristics of OMSW-derived BC

Table 1 shows the effects of pyrolysis temperature on pH, EC, yield, ash, fixed carbon, and volatile matter of OMSW-derived BC. The results showed that BC yield significantly declined with increasing thermal decomposition during the pyrolytic process. The obtained yield of biochar produced at 300 °C accounted for 40.3%, whereas it declined by 32.1%, 28.7%, 27.3%, and 26.7% as the pyrolysis temperature increased to 400, 500, 600, and 700 °C, respectively. This reduction was mainly caused by the organic material decomposition and the dehydration of OH groups during the pyrolysis process. However, increasing thermal decomposition during the pyrolytic process increased the levels of fixed carbon and ash in the OMSW-derived BC. Several other researchers have reported that the pyrolysis temperature is considered one of the main factors determining BC properties, and thus, the effects of its application in the environment [20, 21, 38]. In their studies, the levels of fixed carbon, ash, and basicity increased with increasing temperature of the pyrolytic process, whereas yield and volatile matter of BCs tended to decrease. They suggested that high pyrolysis-temperature BCs possess more carbonaceous materials. Additionally, pyrolysis temperature could affect the thermal stability and chemistry of the BC surface [38].

**Table 1.** pH, electrical conductivity (EC) and approximate analysis of olive mill solid waste-derived biochar (OMSW-BCs) samples as affected by pyrolysis temperature.

| Parameters | FS | BC300 | BC400 | BC500 | BC600 | BC700 |
| --- | --- | --- | --- | --- | --- | --- |
| pH | 5.97 | 9.85 | 10.12 | 10.03 | 10.11 | 10.21 |
| EC, (dS m<sup>-1</sup>) | 0.83 | 1.06 | 2.11 | 2.09 | 2.39 | 2.54 |
| Yield, % | - | 40.3 | 32.1 | 28.7 | 27.3 | 26.7 |
| Moisture, % | 6.11 | 1.16 | 1.44 | 1.72 | 1.89 | 0.95 |
| Volatile matter, % | 74.51 | 27.74 | 17.07 | 11.96 | 11.00 | 9.62 |
| Ash, % | 2.02 | 6.91 | 8.92 | 9.43 | 9.59 | 9.62 |
| Fixed C, % | 17.36 | 64.19 | 72.57 | 76.90 | 77.51 | 79.82 |
| FS: feedstock; BC300: biochar produced at 300 °C; BC400: biochar produced at 400 °C; BC500: biochar produced at 500 °C; BC600: biochar produced at 600 °C; BC700: biochar produced at 700 °C. |  |  |  |  |  |  |

The FTIR results showed that OMSW-derived BC300 possessed a spectrum at 3430 cm^−1^ (Fig. S1), generally attributed to O–H. The spectra of O–H declined with pyrolysis temperature. Additionally, OMSW-derived BC300 exhibited broad bands in the region of 2850–2919 cm^−1^, mainly because of the C-H stretch caused by aliphatic compounds, waxes, and fatty acids. Additionally, the band at 2850 was attributed to symmetrical CH in –CH_2_, suggesting the presence of fatty acids and alkanes. These two peaks at 2850 and 2919 cm-1 declined in BC400 and completely disappeared with a temperature ≥ 500 °C. The absorption at 1574–1652 cm^−1^ of biochar samples suggested the presence of –COOH, as well as C=O and C=C, especially in an aromatic form. The absorption at approximately 1400 cm^−1^ (with an intense band for BC300 and BC400) suggested the presence of aliphatic and aromatic O-H groups.

In this study, XRD was used to identify the mineral composition of BC samples (Fig. S2). In the samples of OMSW-derived BC300, peaks at a spacing of 4.02, 2.06, 1.45, and 1.23 Å were identified, suggesting the presence of kalcinite [K(HCO_3_)], sylvite (KCl), and perclase (MgO), which were reduced or disappeared with increasing pyrolysis temperatures. Furthermore, a high pyrolysis temperature resulted in calcite minerals (CaCO_3_) at 3.04-3.14, 1.87, 1.81 Å. Additionally, the SEM analysis depicted a greater change in the surface structure of BCs compared to that of feedstock with temperature (Fig. S3).

The surface area and pore characteristics of OMSW-derived BCs showed an improvement in the surface area and total and microporous pore volumes with pyrolysis temperature (Table 2). OMSW-derived BCs made at the lowest temperatures of 300 °C and 400 °C, respectively, possessed the lowest surface area of 0.35 m^2^ g^−1^ and 1.78 m^2^ g^−1^; however, OMSW-derived BC500, BC600, and BC700 had high values for the surface area of 108.4, 128.0, and 168.4 m^2^ g^−1^, respectively. In contrast, as pyrolysis temperature increased from 300 °C to 400 °C, 500 °C, 600 °C, and 700 °C, the average pore width for OMSW-derived BCs decreased from to 170.8 to 46.1, 20.4, 19.8, and 19.7 nm, respectively. Generally, during the pyrolytic process, the loss of volatile materials and the functional groups H- and O-were the main reason for the improvement in surface area and pore characteristics of the resultant BCs. In this context, pyrolytic temperatures were positively correlated with BC characteristics, including EC (r = 0.885), fixed carbon (r = 0.9264), surface area (r = 0.9562), microporous surface area (r = 0.9691), total pore volume (r = 0.9532), and microporous pore volume (r = 0.9690) (Table 3). However, pyrolysis temperature showed a negative correlation with yield (r = −0.9054) and volatile matter (r = −9028). In general, the greatest changes were pronounced for high pyrolysis temperatures ≥500 °C. When plotting the biochar characteristics (yield, volatile matter, fixed carbon, ash content, and surface area) in relation to different pyrolysis temperatures, the best model fitting was pronounced with a second-order polynomial (Fig. S4).

**Table 2.** Surface area and pore properties of olive mill solid waste-derived biochars (OMSW-BCs).

| Pyrolysis temperature, °C | SBET<sup>1</sup> (m<sup>2</sup> g<sup>-1</sup>) | S<sub>micr</sub><sup>2</sup> (m<sup>2</sup> g<sup>-1</sup>) | Vt<sup>3</sup> cm<sup>3</sup> g<sup>-1</sup> | V<sub>micro</sub><sup>4</sup> (cm<sup>3</sup> g<sup>-1</sup>) | D<sub>ave</sub><sup>5</sup> (nm) |
| --- | --- | --- | --- | --- | --- |
| 300 | 0.35 | 0.35 | 0.0015 | 0.00018 | 170.8 |
| 400 | 1.78 | 1.78 | 0.0021 | 0.00086 | 46.1 |
| 500 | 108.4 | 80.46 | 0.055 | 0.037 | 20.4 |
| 600 | 128.0 | 109.2 | 0.063 | 0.051 | 19.84 |
| 700 | 168.4 | 143.8 | 0.083 | 0.066 | 19.74 |
| 1. SBET surface area.<br>2. Microporous surface area by the t-plot method.<br>3. Total pore volume<br>4. Microporous pore volume by the t-plot method.<br>5. Average pore width, estimated by 4Vt/SBET |  |  |  |  |  |

**Table 3.**
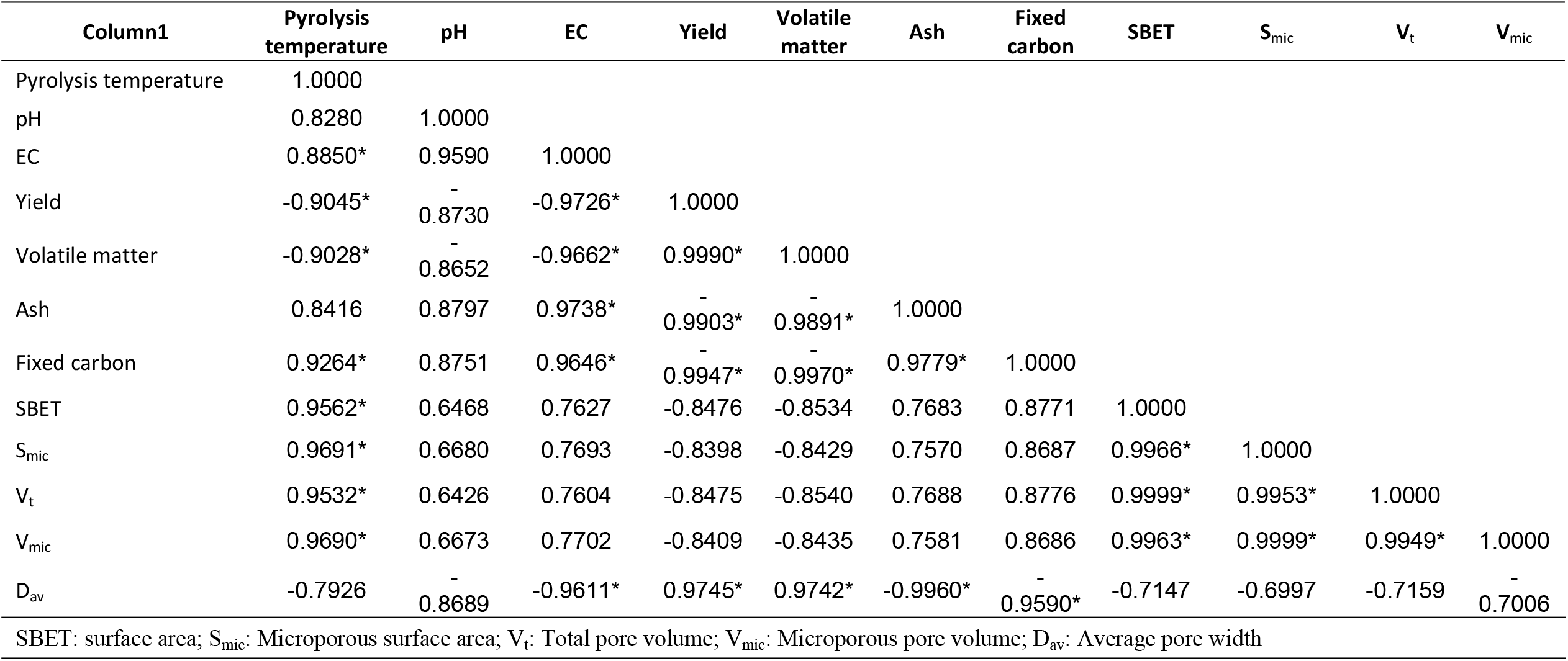
Pearson correlation coefficient (r^2^) among olive mill solid waste-derived biochar (OMSW-BCs) properties.

### Biochar effects on soil pH and EC

OMSW-derived BC treatments of 1% and 3% (w/w) increased soil pH from 7.85 to 8.05 and 8.28 for BC300, 8.15 and 8.31 for BC500, and 8.11 and 8.32 for BC700 (Table 4), mainly because of the alkalinity induced by BCs as a result of the basic oxides and carbonates produced during the pyrolytic process. Previous studies have also indicated the application effects of BCs on increasing soil pH [39–41]. However, in contrast to our results, other studies have found that alkaline soil pH decreased with biochar addition [14, 18]. The reductions in soil pH could be attributed to BC oxidation and the production of acidic materials. In the current study, the higher pH of produced BCs (9.85–10.21) compared to the control soil (7.80) could result in increasing soil pH. The increases in soil pH could have occurred because of the increase in the level of soil exchangeable base cations with BC addition. In a study conducted on calcareous soil, Cardelli et al. [39] suggested that the alkalizing effects of BC on soil pH could be explained by the poor buffering capacity of soil induced by the low levels of soil organic matter.

**Table 4.**
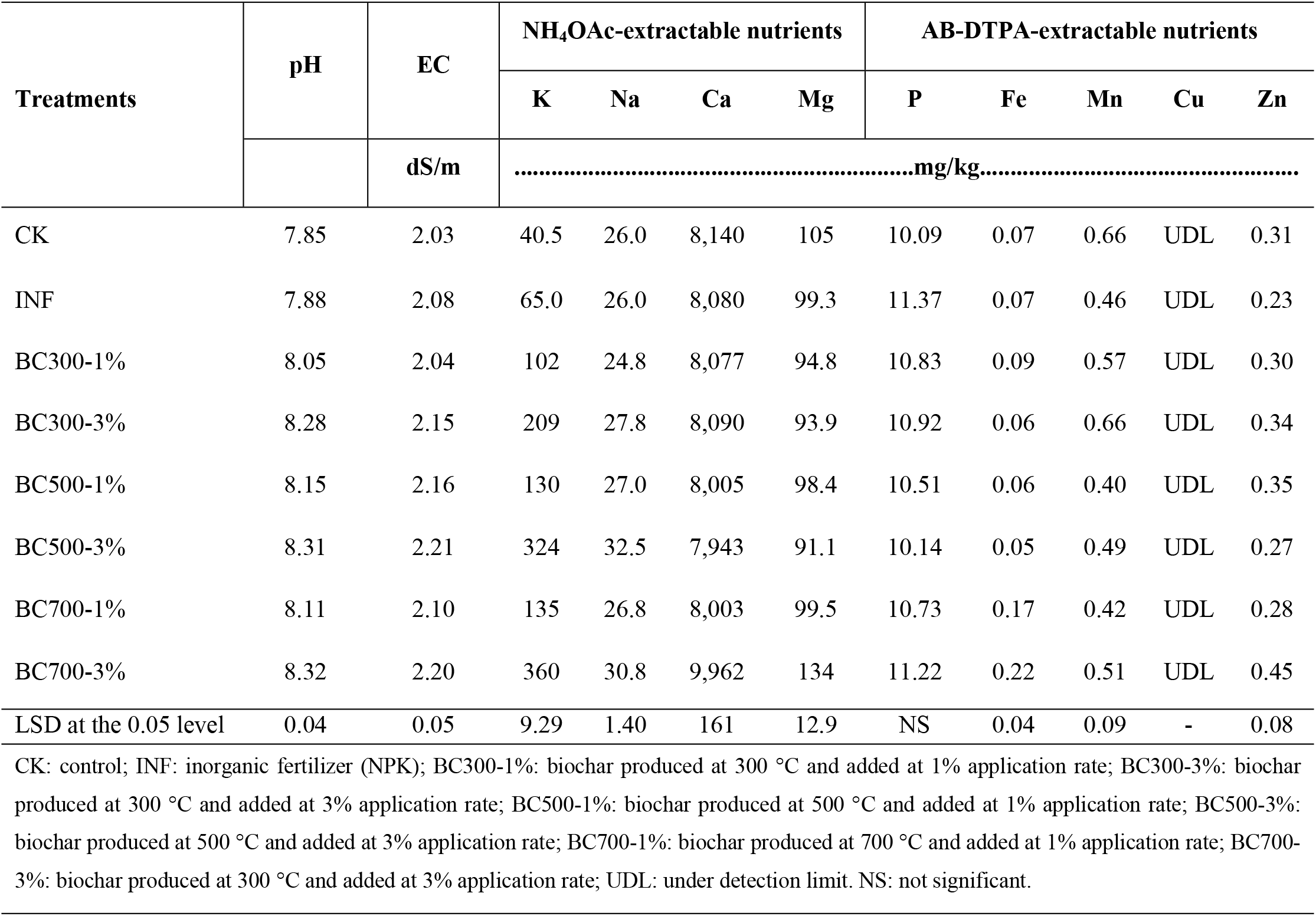
The influences of the applied olive mill solid waste-derived biochars (OMSW-BCs) on the soil pH and EC, and the soil concentrations of NH_4_OAc- and ammonium bicarbonate-diethylenetriaminepentaacetic acid (AB-DTPA)-extractable nutrients.

| Treatments | pH | EC | NH<sub>4</sub>OAc-extractable nutrients |  |  |  | AB-DTPA-extractable nutrients |  |  |  |  |
| --- | --- | --- | --- | --- | --- | --- | --- | --- | --- | --- | --- |
|  |  |  | K | Na | Ca | Mg | P | Fe | Mn | Cu | Zn |
|  |  | dS/m | .....mg/kg..... |  |  |  |  |  |  |  |  |
| CK | 7.85 | 2.03 | 40.5 | 26.0 | 8,140 | 105 | 10.09 | 0.07 | 0.66 | UDL | 0.31 |
| INF | 7.88 | 2.08 | 65.0 | 26.0 | 8,080 | 99.3 | 11.37 | 0.07 | 0.46 | UDL | 0.23 |
| BC300-1% | 8.05 | 2.04 | 102 | 24.8 | 8,077 | 94.8 | 10.83 | 0.09 | 0.57 | UDL | 0.30 |
| BC300-3% | 8.28 | 2.15 | 209 | 27.8 | 8,090 | 93.9 | 10.92 | 0.06 | 0.66 | UDL | 0.34 |
| BC500-1% | 8.15 | 2.16 | 130 | 27.0 | 8,005 | 98.4 | 10.51 | 0.06 | 0.40 | UDL | 0.35 |
| BC500-3% | 8.31 | 2.21 | 324 | 32.5 | 7,943 | 91.1 | 10.14 | 0.05 | 0.49 | UDL | 0.27 |
| BC700-1% | 8.11 | 2.10 | 135 | 26.8 | 8,003 | 99.5 | 10.73 | 0.17 | 0.42 | UDL | 0.28 |
| BC700-3% | 8.32 | 2.20 | 360 | 30.8 | 9,962 | 134 | 11.22 | 0.22 | 0.51 | UDL | 0.45 |
| LSD at the 0.05 level | 0.04 | 0.05 | 9.29 | 1.40 | 161 | 12.9 | NS | 0.04 | 0.09 | - | 0.08 |

CK: control; INF: inorganic fertilizer (NPK); BC300-1%: biochar produced at 300 °C and added at 1% application rate; BC300-3%: biochar produced at 300 °C and added at 3% application rate; BC500-1%: biochar produced at 500 °C and added at 1% application rate; BC500-3%: biochar produced at 500 °C and added at 3% application rate; BC700-1%: biochar produced at 700 °C and added at 1% application rate; BC700-3%: biochar produced at 300 °C and added at 3% application rate; UDL: under detection limit. NS: not significant.

In this study, the EC values for control soils were 2.03 dS m^−1^ (CK) and 2.08 dS m-1 (CK+NPK) (Table 4). These EC values increased to 2.15 dS m-1 for 3% BC300, to 2.16 (BC500 at 1%) and 2.21 dS m-1 (BC500 at 3%), and to 2.10 (BC700 at 1%) and 2.20 dS m-1 (BC700 at 3%). It has been previously reported that a high level of soil salinity following BC application was caused by the introduction of high levels of soluble ions derived from BC ash into the soil [42].

### BC effects on soil nutrient availability

The incorporation of BC into soils can affect soil chemistry, causing changes in the available fraction of soil P, mainly because of alterations in soil pH and cation exchange capacity (CEC). Although soil pH in the current study increased because of the application of OMSW-derived BC, soil AB-DTPA-extractable P concentrations exhibited some increases in the INF and BC treatments compared to that of the control soil, but they were not significant (Table 4). For example, inorganic fertilizer treatment increased soil available P by 1.28 mg kg^−1^, whereas the BC treatments increased its availability by 0.05–1.13 mg kg^−1^. In a meta-analysis study conducted by Glaser and Lehr [16] on the availability of soil P, the addition of BCs to acidic and neutral soils significantly enhanced the availability of soil P, whereas there was no significant effect in alkaline soils. Thus, contrary to the effects in acidic soils, it would be advisable to apply acidic biochar (not alkaline biochar) to alkaline soils with P constraints to improve the levels of plant-available P.

NH_4_OAc-extractable K increased significantly by BC application to the soil (Table 4). In INF-treated soil, the soil exchangeable content of K significantly increased from 40.5 to 65.0 mg kg^−1^. However, because of the application of BCs, the values of soil exchangeable K content greatly increased from 40.5 to 102 mg kg^−1^ (1% BC300), 209 mg kg^−1^ (3% BC300), 130 mg kg^−1^ (1% BC500), 324 mg kg^−1^ (3% BC500), 134 mg kg^−1^ (1% BC700), and 360 mg kg^−1^ (3% BC700). Notably, both pyrolysis temperature and application rates of BCs had a significant effect on the increase in available soil K. Additionally, application of high rates of BCs pyrolyzed at high temperature (500 °C and 700 °C) exhibited greater significant increases in soil exchangeable Na than that in the control soil (Fig. 3), and exhibited increases from 26.0 mg kg^−1^ (CK) to 32.5 mg kg^−1^ (3% BC500) and 30.8 mg kg^−1^ (3% BC700). The experimental results of Jien and Wang [43] indicated that the levels of exchangeable K were significantly enhanced in BC-treated soil, suggesting that it improved the level in the soil. Our results suggested that OMSW-derived BC itself might be a K source, and thus, it enhanced its bioavailability in soils. It has also been reported that BC application could increase the concentration of exchangeable Ca and Mg in the soil [43]. In the present study, among all BC treatments, only 3% BC700 resulted in significant increases in the concentration of NH4OAc-extractable Ca and Mg by 22.4% and 27.9% compared to that of the control soil. This suggested that the concentrations of Ca and Mg in OMSW-derived BC could likely not be available to plants, depending on both pyrolysis temperature and application rate. The high content of soil exchangeable basic cations in the high pyrolysis-temperature BC-treated soil could be explained by the increasing content of BC ash. Several other researchers have explained the improvements in the exchangeability of soil cations because of the presence of ashes in BC, which contain high levels of oxides and hydroxides of alkali cations [44].

Regarding micronutrient availability in soil, BC significantly affected the soil concentrations of AB-DTPA-extractable micronutrients, depending upon pyrolysis temperature and application rates (Table 4). The concentrations of AB-DTPA-extractable Mn in INF, BC500, and BC700 treatments were lower than that in the CK. Although BC treatments caused significant increases in soil pH, application of BC pyrolyzed at the highest temperature (BC700) resulted in a significant improvement in the soil concentrations of AB-DTPA-extractable Fe (at application rates of 1% and 3%) and Zn (at an application rate of 3%). The soil concentrations of AB-DTPA-extractable Fe increased from 0.07 mg kg^−1^ in the CK to 0.17 mg kg^−1^ with 1% BC700 and 0.22 mg kg^−1^ with 3% BC700, whereas AB-DTPA-extractable Zn increased from 0.31 mg kg^−1^ in the CK to 0.45 mg kg^−1^ with 3% BC700. Among all micronutrients, soil concentrations of AB-DTPA-extractable Cu were undetectable by ICP-OES.

### BC effects on dry matter and mineral content of maize plants

Statistically, BC addition did not significantly affect plant growth parameters (fresh and dry weight and plant height) (Table 5). Similarly, Farrell et al. [45] (2014) found no improvement in wheat yield in calcareous soils because of BC addition. They suggested that the addition of BC to calcareous soil did not lead to better conditions for the uptake of plant nutrients. In this context, the results of the current study showed that the shoots of maize plants in BC-amended soils had significantly lower content of P, Ca, and Mg. In soil treated with 3% OMSW-derived BC, the shoot P content decreased from 2.24 g kg^−1^ (CK) to 1.68 g kg^−1^ (BC300), 1.82 g kg^−1^ (BC500), and 1.88 g kg^−1^ (BC700). The shoot content of Ca decreased from 13.06 g kg^−1^ in the CK to 10.50, 7.64, and 6.23 g kg^−1^ for BC300, BC500, and BC700, respectively. Furthermore, the shoot content of Mg decreased from 10.6 g kg^−1^ in the CK to 5.08, 3.88, and 3.71 g kg^−1^ for BC300, BC500, and BC700, respectively. In alkaline soils, calcareous substances can lead to the formation of insoluble compounds of Ca-Mg-P, decreasing the shoot content of these nutrients [46]. On the other hand, the great quantity of K obtained because of OMSW-derived BC could result in reduced plant uptake of Ca, Mg, and P by an antagonistic effect [47, 48]. Contrary to our results, several reports found increases in the plant content of nutrients [44, 49]. The discrepancy between our data and that of other studies could be explained because of varying soil characteristics and feedstock used to produce BCs. Our results suggested that the incorporation of OMSW-derived BCs into alkaline soils may limit the essential nutrient (such as Ca, Mg, and P) uptake by plants, and require the additional input of these nutrients (especially P fertilizer). Further research on OMSW-derived BCs is needed to clarify the mechanism responsible for governing uptake of these nutrients by plants.

**Table 5.** Olive mill solid waste-derived biochar (OMSW-BCs) effects on plant height, fresh and dry matter, and shoot mineral content of *Zea mays*.

| Treatments | Plant height (cm) | Plant fresh weight (g plant<sup>-1</sup>) | Plant dry (g plant<sup>-1</sup>) | K | Na | P | Ca | Mg | Fe | Mn | Zn | Cu |
| --- | --- | --- | --- | --- | --- | --- | --- | --- | --- | --- | --- | --- |
|  |  |  |  | g kg<sup>-1</sup> |  |  |  |  | mg kg<sup>-1</sup> |  |  |  |
| CK | 54.5 | 2.37 | 0.21 | 15.1 | 0.14 | 2.24 | 13.06 | 10.6 | 108 | 49.5 | 64.5 | 8.94 |
| INF | 54.3 | 2.56 | 0.22 | 23.4 | 0.16 | 2.41 | 16.60 | 9.51 | 106 | 53.9 | 76.3 | 8.22 |
| BC300-1% | 54.2 | 2.59 | 0.20 | 33.2 | 0.26 | 1.71 | 12.37 | 6.45 | 91.3 | 43.1 | 74.3 | 7.39 |
| BC300-3% | 50.2 | 2.58 | 0.18 | 56.1 | 0.30 | 1.68 | 10.50 | 5.08 | 105 | 39.4 | 71.9 | 9.67 |
| BC500-1% | 53.6 | 2.76 | 0.21 | 49.9 | 0.28 | 2.18 | 11.61 | 5.90 | 105 | 47.1 | 81.8 | 8.11 |
| BC500-3% | 49.7 | 2.53 | 0.20 | 65.3 | 0.31 | 1.82 | 7.64 | 3.88 | 92.4 | 43.1 | 79.3 | 7.56 |
| BC700-1% | 59.4 | 3.17 | 0.24 | 39.4 | 0.51 | 2.17 | 11.35 | 6.33 | 108 | 46.2 | 77.4 | 7.72 |
| BC700-3% | 49.4 | 2.62 | 0.21 | 73.7 | 0.79 | 1.88 | 6.23 | 3.71 | 97.7 | 40.2 | 69.2 | 7.22 |
| LSD at the | NS | NS | NS | 10.94 | 0.05 | 0.26 | 1.94 | 0.75 | NS | 5.28 | 12.68 | NS |

CK: control; INF: inorganic fertilizer (NPK); BC300-1%: biochar produced at 300 °C and added at 1% application rate; BC300-3%: biochar produced at 300 °C and added at 3% application rate; BC500-1%: biochar produced at 500 °C and added at 1% application rate; BC500-3%: biochar produced at 500 °C and added at 3% application rate; BC700-1%: biochar produced at 700 °C and added at 1% application rate; BC700-3%: biochar produced at 300 °C and added at 3% application rate; NS: not significant.

Contrary to that of shoot P, Ca, and Mg, OMSW-derived BCs treatments enhanced the levels of K and Na in plant shoots (Table 5). In soil treated with 1% and 3% OMSW-derived BC, respectively, the shoot K content increased from 15.1 g kg^−1^ (CK) and 23.4 g kg^−1^ (INF) to 33.2 and 56.1 g kg^−1^ (BC300), 49.9 and 65.3 g kg^−1^ (BC500), and 39.4 and 73.7 g kg^−1^ (BC700). This indicated that shoot K content increased with increasing pyrolysis temperature and application rate. In this study, soil exchangeable K was also significantly increased following BC application and it increased with both increasing pyrolysis temperature and application rate, reflecting enhanced K uptake by plant shoots. It has been reported that a high quantity of K in BCs could enhance the bioavailable pool of K in the soil [50]. Application of BCs might also improve soil mineral K release by enhancing the activity of K-dissolving bacteria and facilitating K uptake by crops [51]. Syuhada et al. [52] found that the K content in corn tissue was significantly higher but its tissue content of N, Ca, and Mg was lower in BC-amended plants than that of the control. They attributed the high tissue K content in BC treatments to increasing exchangeable soil K.

BCs exhibited varying effects on the shoot content of micronutrients, depending upon the micronutrient. Statistically, BC addition did not have a significant effect on the shoot levels of Fe and Cu (Table 5). However, the addition of BCs (especially at the highest application rate) significantly decreased the shoot Mn levels in comparison with that of the CK. Conversely, application of 1% and 3% BC500 and 1% BC700 significantly enhanced the level of shoot Zn.

### Recommendations and suggestions for future trials

This study was conducted to evaluate the effects of application of OMSW-derived BCs (in relation to pyrolysis temperature and application rates) on soil pH, EC, availability of soil nutrients to plants, and maize growth in arid alkaline soil. The high K content in BC treatments suggests high agronomic value in terms of replacing conventional K sources. However, despite the high soil available concentrations of K, the significant increases in soil pH values and decreases in P, Ca, Mg, and Mn of plant tissues because of the application of BC should not be ignored because these nutrients are of importance in terms of agronomic potential. The quantity of K incorporated by OMSW-derived BC could negatively reflect the decreasing nutrient uptake by plants, as indicated in this study for the decrease in shoot content of P, Ca, Mg, and Mn. For OMSW-derived BCs, lowering of shoot nutrient content may likely limit its use for BC production, and thus, care should be taken in its use under arid conditions. However, because of its high alkalinity, it could be speculated that OMSW-derived BCs may be applied as a soil additive to acidic soils rather than alkaline ones. Further studies on OMSW-derived BCs are needed to evaluate its effects on a wide range of alkaline and calcareous soils with varied properties. Further studies are also required on the effects of OMSW-derived BC in combination with inorganic fertilizers on the antagonistic effect induced by the high quantity of K and the properties and fertility of arid-land soils.

### Conclusions

This study was established to investigate (i) pyrolysis temperature (300–700 °C) effects on the properties of OMSW-derived BCs, and (ii) short-term application effects of OMSW-derived BCs on pH, EC, and nutrients availability in calcareous sandy soil, as well as its effects on maize plant growth. An increase in pyrolytic temperature resulted in increased biochar pH, EC, ash content, fixed carbon content, and surface area but decreased yield and volatile matter content. The results also showed that OMSW-derived BC increased soil pH and EC and enhanced the levels of soil available K, Na, Fe, and Zn, primarily depending on the pyrolysis temperature and application rate. The application of OMSW-derived BC decreased the levels of Ca, Mg, Mn, and P in plant shoots and enhanced the levels of shoots K, Na, and Zn. Thus, it can be concluded that application of OMSW-derived BCs as soil additives may enhance the availability of soil K, as well as its content in plant tissues; nonetheless, it may cause negative effects because of decreasing levels of essential nutrients in plant tissues. Collectively, our findings suggest that application of OMSW-derived BCs may not be able to provide sufficient nutrients to enhance plant growth.

## Acknowledgments

The authors extend their appreciation to the Deanship of Scientific Research, King Saud University, for funding this work through the international research group project RG-1439-043.

## Author contribution

**Conceptualization:** MIA; ARAU.

**Data curation:** ARAU; ASS; MA.

**Formal analysis:** AA; MAA; JE.

**Funding acquisition:** MIA.

**Investigation:** AA; MAA; HI; JE.

**Methodology:** AA; MAA.

**Project administration:** MIA.

**Resources:** MIA.

**Supervision:** MIA; ARAU.

**Validation:** ARAU; AA.

**Visualization:** MIA; AA; ARAU.

**Writing - original draft:** ARAU; AA.

**Writing - review & editing:** ARAU; MIA; AA.

## Supporting Information

Fig. S1. FTIR spectra of the produced olive mill solid waste-derived biochar (OMSW-BCs) (BC300: biochar produced at 300 °C; BC400: biochar produced at 400 °C; BC500: biochar produced at 500 °C; BC600: biochar produced at 600 °C; BC700: biochar produced at 700 °C).

Fig.S2. XRD spectra of the produced olive mill solid waste-derived biochar (OMSW-BCs) (BC300: biochar produced at 300 °C; BC400: biochar produced at 400 °C; BC500: biochar produced at 500 °C; BC600: biochar produced at 600 °C; BC700: biochar produced at 700 °C).

Fig. S3. Scanning electron microscope (SEM) analyses of feedstock (FS) and olive mill solid waste-derived biochars (OMSW-BCs) pyrolyzed at different temperatures (a: FS: feedstock; b: BC300: biochar produced at 300 °C; c: BC400: biochar produced at 400 °C; d: BC500: biochar produced at 500 °C; e: BC600: biochar produced at 600 °C; f: BC700: biochar produced at 700 °C).

Fig. S4. Model fitting for the produced olive mill solid waste-derived biochar (OMSW-BCs) properties in relation with pyrolysis temperatures.

## References

1. Garibaldi LA, Gemmill-Herren B, D’Annolfo R, Graeub BE, Cunningham SA, Breeze TD. Farming approaches for greater biodiversity, livelihoods, and food security. Trends Ecol Evol. 2017; 32(1): 68–80. https://doi.org/10.1016/j.tree.2016.10.001.

2. Biswas S, Ali MN, Goswami R, Chakraborty S. Soil health sustainability and organic farming: A review. J Food Agri. Environ. 2014; 12(3&4): 237–243.

3. Rajper AM, Udawatta RP, Kremer RJ, Lin C, Jose S. Effects of probiotics on soil microbial activity, biomass and enzymatic activity under cover crops in field and greenhouse studies Agrofor Syst. 2016; 90: 811–827. https://doi.org/10.1007/s10457-016-9895-1.

4. McGeehan SL. Impact of waste materials and organic amendments on soil properties and vegetative performance. Appl Environ Soil Sci. 2012; Volume 2012, Article ID 907831, 11 pages. https://doi.org/10.1155/2012/907831.

5. Doan TT, Henry-des-Tureaux T, Rumpel C, Janeau JL, Jouquet P. Impact of compost, vermicompost and biochar on soil fertility, maize yield and soil erosion in Northern Vietnam: a three-year mesocosm experiment. Sci Total Environ. 2015; 514:147–154. https://doi.org/10.1016/j.scitotenv.2015.02.005.

6. Li L-J, Han X-Z. Changes of soil properties and carbon fractions after long-term application of organic amendments in Mollisols. CATENA. 2016; 143:140–144. https://doi.org/10.1016/j.catena.2016.04.007.

7. Luo G, Li L, Friman, V-P, Guo J, Guo S, Shen Q, et al. Organic amendments increase crop yields by improving microbe-mediated soil functioning of agroecosystems: A meta-analysis. Soil Biol Biochem. 2018; 124:105–115. https://doi.org/10.1016/j.soilbio.2018.06.002.

8. Wu Y, Xu G, Shao HB. Furfural and its biochar improve the general properties of a saline soil. Solid Earth. 2014: 5:665–671. https://doi.org/10.5194/se-5-665-2014.

9. Ge X, Cao Y, Zhou B, Wang X, Yang Z, Li M-H. Biochar addition increases subsurface soil microbial biomass but has limited effects on soil CO2 emissions in subtropical moso bamboo plantations. Appl Soil Ecol. 2019; 142:155–165. https://doi.org/10.1016/j.apsoil.2019.04.021.

10. Yu H, Zoucd W, Chen J, Chen H, Yu Z, Huang J, et al. Biochar amendment improves crop production in problem soils: A review. J Environ Manage. 232: 8–21. https://doi.org/10.1016/j.jenvman.2018.10.117.

11. Yaashikaa PR, Kumar P.S., Varjani, S. J., and Saravanan. A.: Advances in production and application of biochar from lignocellulosic feedstocks for remediation of environmental pollutants. Bioresour Technol. 2019; 292:122030. https://doi.org/10.1016/j.biortech.2019.122030.

12. Lehmann J, da Silva JP, Steiner C, Nehls T, Zech W, Glaser B. Nutrient availability and leaching in an archaeological Anthrosol and a Ferralsol of the Central Amazon basin: fertiliser, manure and charcoal amendments. Plant Soil. 249:343–357. https://doi.org/10.1023/A:102283311.

13. Baiamonte G, Crescimanno G, Parrino F, De Pasquale C. Effect of biochar on the physical and structural properties of a sandy soil. CATENA, 2019; 175:294–303. https://doi.org/10.1016/j.catena.2018.12.019.

14. Zhang M, Riaz M, Zhang L, El-desouki Z, Jiang. C. Biochar induces changes to basic soil properties and bacterial communities of different soils to varying degrees at 25 mm rainfall: More effective on acidic soils. Front Microbiol. 2019: 10:1321. https://doi:10.3389/fmicb.2019.01321.

15. Dai Z, Wang Y, Muhammad N, Yu X, Xiao K, Meng J, et al. The effects and mechanisms of soil acidity changes, following incorporation of biochars in three soils differing in initial pH. Soil Sci Soc Am J. 2014; 78:1606–1614. https://doi:10.2136/sssaj2013.08.0340.

16. Glaser B, Lehr V-I. Biochar effects on phosphorus availability in agricultural soils: A meta-analysis. Sci Rep. 2019; 9: 9338. https://doi.org/10.1038/s41598-019-45693-z.

17. Liu XH, Zhang XC. Effect of biochar on pH of alkaline soils in the Loess Plateau: Results from incubation experiments. Int. J. Agric. Biol. 2012; 4:745–750.

18. Abrishamkesh, S, Gorji, M, Asadi, H, Bagheri-Marandi, GH, Pourbabaee, AA. Effects of rice husk biochar application on the properties of alkaline soil and lentil growth. Plant Soil Environ. 2015; 61(11), 475–482. https://doi.org/10.17221/117/2015-PSE.

19. Salem TM, Refaie KM, Sherif AEEAE, Eid MAM. Biochar application in alkaline soil and its effect on soil and plant. Acta Agric Slov. 2019; 114:85–96. https://doi:10.14720/aas.2019.114.1.10.

20. Usman AR, Abduljabbar A, Vithanage M, Ok YS, Ahmad M, Ahmad M, Elfakia J, et al. Biochar production from date palm waste: Charring temperature induced changes in composition and surface chemistry. J Anal Appl Pyrolysis. 2015; 115:392–400. https://doi.org/10.1016/j.jaap.2015.08.016.

21. Li S, Harris S, Anandhi A, Chen G. Predicting biochar properties and functions based on feedstock and pyrolysis temperature: A review and data syntheses. J Clean Prod. 2109; 215:890–902. https://doi.org/10.1016/j.jclepro.2019.01.106.

22. Al-Wabel MI, Al-Omran A, El-Naggar AH, Nadeem, Usman AR. Pyrolysis temperature induced changes in characteristics and chemical composition of biochar produced from conocarpus wastes. Bioresour Technol. 2013; 131:374–379. https://doi.org/10.1016/j.biortech.2012.12.165.

23. Roig A, Cayuela, ML, Sanchez-Monedero MA. An overview on olive mill wastes and their valorisation methods. Waste Manage. 2006; 26(9): 960–969. https://doi.org/10.1016/j.wasman.2005.07.024.

24. Arjona R, Ollero P, Vidal F. Automation of an olive waste industrial rotary dryer. J Food Eng. 2005; 68:239–247. https://doi:10.1016/j.jfoodeng.2004.05.049.

25. Obied HK, Allen MS, Bedgood DR, Prenzler PD, Robards K, Stockmann R. Bioactivity and analysis of biophenols recovered from olive mill waste. J Agric Food Chem. 2005; 53:823–837. https://doi:10.1021/jf048569x.

26. Hmid A, Al Chami Z, Sillen W, De Vocht A, Vangronsveld J. Olive mill waste biochar: a promising soil amendment for metal immobilization in contaminated soils. Environ Sci Pollut Res Int. 2015; 22(2): 444–1456. https://doi:10.1007/s11356-014-3467-6.

27. Cayuela ML, Bernal MP, Roig A. Composting olive mill wastes and sheep manure for orchard use. Compost Sci Util. 2004; 12(2):130–136. https://doi.org/10.1080/1065657X.2004.10702171.

28. Alcaide EM, Nefzaoui A. Recycling of olive oil byproducts: Possibilities of utilization in animal nutrition. Int Biodeterior Biodegradation. 1996; 38(3/4):227–235. https://doi.org/10.1016/S0964-8305(96)00055-8.

29. Abdelhadi SO, Dosoretz CG, Giora R, Gerchman Y and Azaizeh, H. Production of biochar from olive mill solid waste for heavy metal removal. Bioresour Technol. 2015; 244:759–767. https://doi.org/10.1016/j.biortech.2017.08.013.

30. Piscitelli L, Shaaban A, Mondelli D, Mezzapesa GN, Miano TM, Dumontet, S. Use of olive mill pomace biochar as a support for soil microbial communities in an Italian sandy soil. Soil Horizons. 2015; 56:1–7. https://doi:10.2136/sh15-02-0006.

31. Senbayram M, Saygan EP, Chen R, Aydemir S, Kaya C, Wu D, Bladogatskaya, E. Effect of biochar origin and soil type on the greenhouse gas emission and the bacterial community structure in N fertilised acidic sandy and alkaline clay soil. Sci Total Environ. 2019; 660:69–79. https://doi:10.1016/j.scitotenv.2018.12.300.

32. ASTM D1762-84. Standard methods for chemical analysis of wood charcoal, ASTM International, West Conshohocken, PA, USA, 2013.

33. Bouyoucos, G. J. Hydrometer method improved for making particle size analysis of soils, J. Argon. 1962; 53:464–465. https://doi:10.2134/agronj1962.00021962005400050028x.

34. Sparks, D L. 1996. Methods of soil analysis. Part 3. Chemical methods. Am. Soc. of Agron., Inc. Madison, WI., 1996.

35. Nelson, D. W., and Sommers, L.E. Total carbon, organic carbon, and organic matter. In Methods of Soil Analysis. Part 3. Chemical Methods. Edited by Sparks et al., SSSA and ASA, Madison, WI. pp. 961–1010, 1996.

36. Lambert MJ. Preparation of plant material for estimating a wide range of elements. Forestry Commission of New South Wales, Sydney, 29, 1–60, 1976.

37. StatSoft, Inc. 1995. Statistica for Windows (Computer Program Manual). StatSoft, Inc., Tulsa, OK., 1995.

38. Tan C, Yaxin Z, Hongtao W, Wenjing L, Zeyu Z, Yuancheng Z. et al. Influence of pyrolysis temperature on characteristics and heavy metal adsorptive performance of biochar derived from municipal sewage. Bioresour. Technol. 2014; 164:47–54. https://doi:10.1016/j.biortech.2014.04.048.

39. Cardelli R, Becagli M, Marchini F, Saviozzi A. Effect of biochar, green compost, and vermicompost on the quality of a calcareous soil: A 1-Year laboratory experiment. Soil Sci. 2017; 182:1–8. https://doi:10.1097/SS.0000000000000216.

40. Yao Q, Liu J, Yu Z, Li Y, Jin J, Liu X. et al. Three years of biochar amendment alters soil physiochemical properties and fungal community composition in a black soil of northeast China. Soil Biol Biochem. 2017; 110:56–67. https://doi.org/10.1016/j.soilbio.2017.03.005.

41. Tang, J, Cao C, Gao F, Wang W. Effects of biochar amendment on the availability of trace elements and the properties of dissolved organic matter in contaminated soils. Environ. Technol. Inno. 2019; 16:100492. https://doi.org/10.1016/j.eti.2019.100492.

42. Chintala R, Mollinedo J, Schumacher TE, Malo DD, Julson JL. Effect of biochar on chemical properties of acidic soil. Arch. Agron. Soil Sci. 2014; 60:393–404. https://doi.org/10.1080/03650340.2013.789870.

43. Jien S-H, Wang C-S. Effects of biochar on soil properties and erosion potential in a highly weathered soil. CATENA. 2013; 110:225–233. https://doi.org/10.1016/j.catena.2013.06.021.

44. da Silva ICB, Basílio, JJN, Fernandes LA, Colen F, Sampaio RA, Frazão LA. Biochar from different residues on soil properties and common bean production. Sci. Agric. 74(5):378–382. https://doi.org/10.1590/1678-992x-2016-0242.

45. Farrell M, Macdonald LM, Butler G, Chirino-Valle I, Condron LM. Biochar and fertiliser applications influence phosphorus fractionation and wheat yield. Biol Fertil Soils. 2014; 50:169. https://doi.org/10.1007/s00374-013-0845-z.

46. Chintala R, Schumacher TE, McDonald LM, David DE, Clay E, Malo DD, et al. Phosphorus sorption and availability from biochars and soil/biochar mixtures. Clean Soil, Air, Water. 2014; 42:626–634. https://doi.org/10.1002/clen.201300089.

47. Jakobsen ST. Interaction between Plant Nutrients: III. Antagonism between potassium, magnesium and calcium. Acta Agr Scand. B Soil Plant Sci. 1993; 43:1–5. https://doi.org/10.1080/09064719309410223.

48. Li Z, Zhang R, Xia S, Wang L, Liu C, Zhang R, et al. Interactions between N, P and K fertilizers affect the environment and the yield and quality of satsumas. Glob Ecol Conserv. 2019; 19, e00663. https://doi.org/10.1016/j.gecco.2019.e00663.

49. Shen Q, Hedley M, Arbestain MC, Kirschbaum MUF. Can biochar increase the bioavailability of phosphorus?,. J Soil Sci Plant Nutr. 2016; 16(2):268–286. https://doi:10.4067/S0718-95162016005000022.

50. Tian X, Li C, Zhang M, Wan Y, Xie Z, Chen B, et al. Biochar derived from corn straw affected availability and distribution of soil nutrients and cotton yield. PLoS ONE. 2018; 13(1): e0189924. https://doi.org/10.1371/journal.pone.0189924.

51. Wang L, Xue C, Nie X, Liu Y, Chen F. Effects of biochar application on soil potassium dynamics and crop uptake. J Plant Nutr. Soil Sci. 2018; 181:635–643. https://doi.org/10.1002/jpln.201700528.

52. Syuhada AB, Shamshuddin J, Fauziah CI, Rosenani, AB, Arifin A. Biochar as soil amendment: Impact on chemical properties and corn nutrient uptake in a Podzol. Can J Soil Sci. 2016; 96(4): 400–412. https://doi.org/10.1139/cjss-2015-0044.

